# ICA512 RESP18 homology domain is a protein condensing factor and insulin fibrillation inhibitor

**DOI:** 10.1101/521351

**Authors:** Pamela L. Toledo, Juha M. Torkko, Andreas Müller, Carolin Wegbrod, Anke Sönmez, Michele Solimena, Mario R. Ermácora

**Affiliations:** Grupo de Biología Estructural y Biotecnología, IMBICE, CONICET, Universidad Nacional de Quilmes, Bernal, Buenos Aires, Argentina; Dept. Molecular Diabetology, University Hospital and Faculty of Medicine, TU Dresden, Dresden, Germany; Paul Langerhans Institute Dresden of the Helmholtz Center Munich at the University Hospital and Faculty of Medicine, TU Dresden, Dresden, Germany; German Center for Diabetes Research (DZD e.V.), Neuherberg, Germany; Dept. Molecular Diabetology, University Hospital and Faculty of Medicine, TU Dresden, Dresden, Germany; Paul Langerhans Institute Dresden of the Helmholtz Center Munich at the University Hospital and Faculty of Medicine, TU Dresden, Dresden, Germany; German Center for Diabetes Research (DZD e.V.), Neuherberg, Germany; Max Planck Institute of Molecular Cell Biology and Genetics, Dresden, Germany

**Keywords:** ICA512, IA-2, PTPRN, insulin, β-cell, protein aggregation, secretory granule, secretion, protein targeting, diabetes

## Abstract

Type 1 diabetes islet cell autoantigen 512 (ICA512) is a tyrosine phosphatase-like intrinsic membrane protein involved in the biogenesis and turnover of insulin secretory granules (SGs) in pancreatic islet β-cells. Whereas its membrane proximal and cytoplasmic domains have been functionally and structurally characterized, the role of ICA512 N-terminal segment named ‘regulated endocrine-specific protein 18 homology domain’ (RESP18HD), which encompasses residues 35–131, remains largely unknown. Here we show that ICA512 RESP18HD residues 91–131 encode for an intrinsically disordered region (IDR), which *in vitro* acts as a condensing factor for the reversible aggregation of insulin and other β-cell proteins in a pH and Zn^2+^regulated fashion. At variance with what has been shown for other granule cargoes with aggregating properties, the condensing activity of ICA512 RESP18HD is displayed at pH close to neutral, *i.e*. in the pH range found in the early secretory pathway, while it is resolved at acidic pH and Zn^2+^ concentrations resembling those present in mature SGs. Moreover, we show that ICA512 RESP18HD residues 35–90, preceding the IDR, inhibit insulin fibrillation *in vitro*. Finally, we found that glucose-stimulated secretion of RESP18HD upon exocytosis of SGs from insulinoma INS-1 cells is associated with cleavage of its IDR, conceivably to prevent its aggregation upon exposure to neutral pH in the extracellular milieu. Taken together, these findings point to ICA512 RESP18HD being a condensing factor for protein sorting and granulogenesis early in the secretory pathway, and for prevention of amyloidogenesis.

## Introduction

Receptor-type protein tyrosine phosphatases (RPTPs) are transmembrane proteins involved in signaling pathways (1). ICA512 (also known as IA-2, PTP35, or PTPRN) and phogrin (also known as IA-2β, IAR, ICAAR, or PTPRN2) are R8 subtype RPTPs mainly expressed in peptide-hormone secreting endocrine cells and neurons, where they are enriched in the membrane of secretory granules (SGs) and large dense-core vesicles, respectively (2, 3).

In mice, genetic deletion of ICA512, phogrin, or both results in mild glucose intolerance and decreased glucose-responsive insulin secretion (4–6). A large body of evidence suggests that both proteins are involved in the biogenesis and turnover of insulin SGs in pancreatic islet β-cells (7–9). Accordingly, their depletion, either alone or in combination, strongly reduces – albeit does not abolish– insulin SG stores *in vitro* and *in vivo*.

ICA512 and phogrin have a large lumenal ectodomain, a single-pass transmembrane segment, and a cytoplasmic catalytically-impaired, protein tyrosine phosphatase (PTP) domain (Fig. 1*A*) (10). During SG maturation, ICA512 ectodomain is cleaved by furin-like convertases, generating a N-terminal fragment (ICA512 NTF; residues 35–448) and a transmembrane fragment (ICA512 TMF; residues 449–979) (3).

**Fig. 1.**
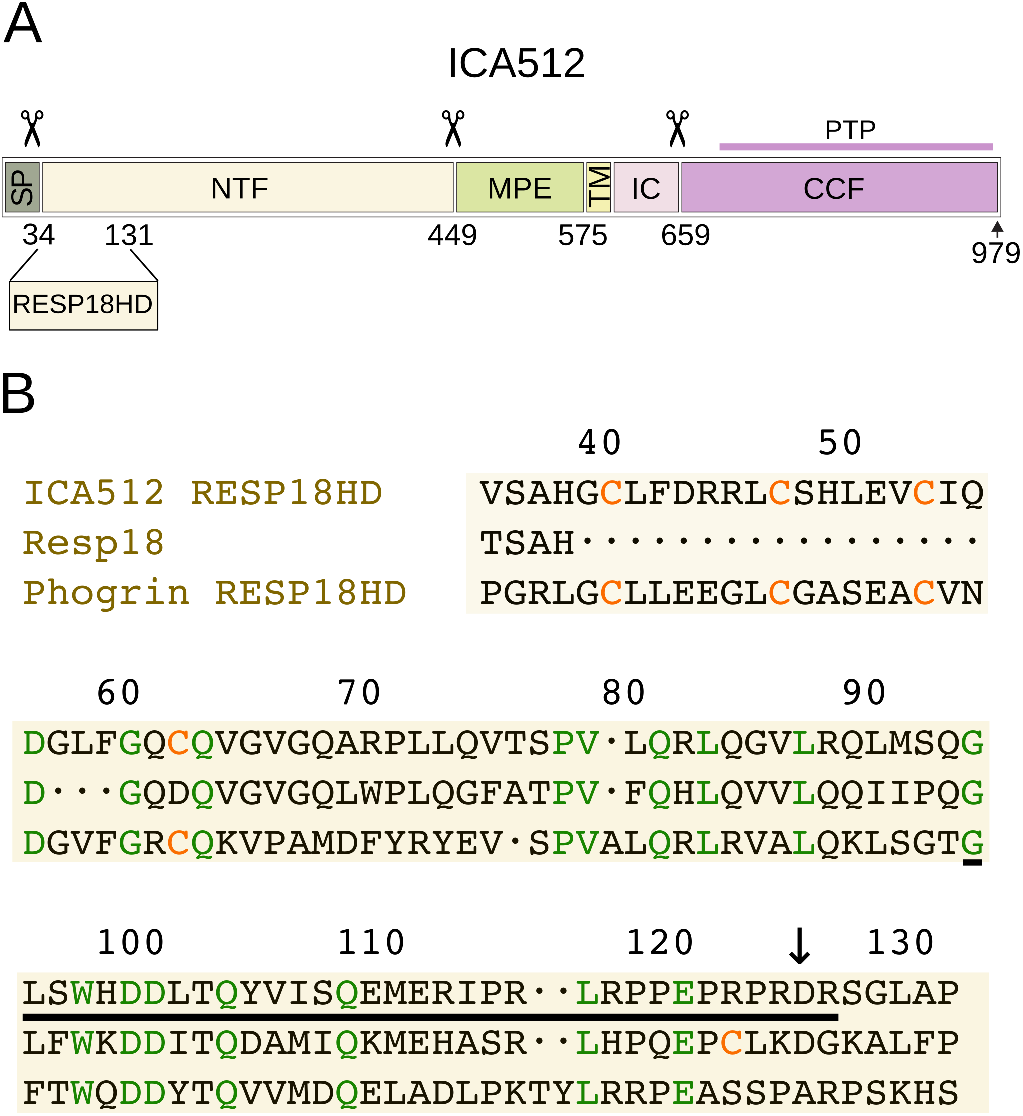
ICA512 processing and ICA512 RESP18HD sequence. *A*: the SG lumenal segment of ICA12 (UniProtKB Q16849) comprises residues 1–575 and includes a signal peptide (*SP*), the N-terminal fragment (*NTF*), and the Membrane Proximal Ectodomain (*MPE*). The transmembrane domain (*TM*) comprises residues 576–600. The cytoplasmic region, residues 601–979, is made of the jux-tamembrane intracellular domain (*IC*) and the Cytoplasmic Cleaved Fragment (*CCF*). Most of *CCF* corresponds to the pseudo-phosphatase (catalytically inactive) domain (*PTP*), from which the entire protein is named. Scissors mark well characterized processing sites. Convertase mediated cleavage at residue 448 generates the transmembrane fragment (ICA512 TMF, residues 449–979, corresponding to MPE–TM–IC), which is transiently inserted into the plasma membrane upon exocytosis. ICA512 RESP18HD constitutes the N-terminal portion of ICA512 NTF. *B*: sequence alignment of ICA512 RESP18HD, Resp18 (UniProtKB Q5W5W9), and phogrin RESP18HD (UniProtKB Q92932). Identical residues and cysteine residues are indicated in green and orange, respectively. An ICA512 RESP18HD C-terminal cleavage site between R124 and D125 in ICA512-RESP18HD (this manuscript) is indicated with a down arrow. Most of the IDR accounting for ICA512 RESP18HD aggregation activity (this manuscript) is the product of exon-4, which encodes for ICA512 residues 94–126 (underlined).

The cytoplasmic domain of ICA512 TMF interacts with the cortical cytoskeleton via β-2-syntrophin, an adapter protein involved in the tethering of SGs to F-actin, thereby regulating insulin SG mobility and exocytosis (9, 11, 12). Furthermore, through an still unidentified signalling pathway, ICA512 modulates gene expression of the F-actin modifier villin, another factor for tight control of insulin SG mobility and exocytosis (13).

Transient insertion of ICA512 TMF in the plasma membrane upon insulin SG exocytosis is coupled to the cleavage of its cytoplasmic tail by Ca^2+^-dependent calpain (12) and the generation of a cytoplasmic fragment that enhances STAT5-mediated transcription of SG cargoes and mouse β-cell proliferation (8, 11, 14, 15).

The SG lumenal, *N*-glycosylated juxtamembrane region of ICA512 TMF comprises the membrane-proximal ectodomain (ICA512 MPE). X-ray crystallography revealed that ICA512 MPE is a SEA (**S**ea urchin sperm protein, **E**nterokinase, **A**grin) domain, which is compatible with its potential involvement in cell adhesion and the formation of homo- and heterodimers (16–19). Further studies indicated that β-strands in ICA512 MPE promote ICA512 dimerization and regulate the exit of the proprotein from the endoplasmic reticulum (20). A similar SEA domain is also found in the corresponding region of phogrin (21, 22).

ICA512 NTF –the other lumenal segment resulting from the proprotein convertase processing of ICA512– is much less characterized. The ICA512 NTF region encompassing residues 35–131, immediately after the signal peptide, exhibits sequence similarity to the regulated endocrine-specific protein 18 (Resp18; Fig. 1). As ICA512 and phogrin, Resp18 is widely expressed in peptide secreting cells, including α, β and δ cells of the pancreatic islets and its expression is up regulated in conditions stimulating SG biogenesis (23).

In view of its similarity to Resp18, the segment corresponding to residues 35–131 of ICA512 was named Resp18 homology domain (RESP18HD) (24). Interestingly, ICA512 RESP18HD has a conserved cysteine-rich N-terminal region (residues 35–62), which is absent in Resp18 (Fig. 1*B*.

In previous studies, we showed that ICA512 RESP18HD contains sufficient information to direct green fluorescent protein (GFP) to insulin SGs, whereas deletion of the entire ICA512 NTF, which includes ICA512 RESP18HD, causes the constitutive delivery of ICA512 to the plasma membrane, hence abolishing SG targeting (20, 24).

While the detailed mechanism of proprotein sorting into SGs is not well established, it is clear that signal-mediated sorting and congregation–aggregation of regulated cargoes contribute to the biogenesis of SGs (25–28).

Congregation–aggregation processes are of two main classes: liquid–liquid phase transitions (LLPT) and liquid–solid phase separation or, for short, aggregation. LLPT and aggregation have in common a stage of segregation in which the participant molecules congregate guided by multiple non-covalent interactions. However, the outcomes of these two processes are different, being either separated liquid-like or solid-like phases. These two kinds of phases establish a dynamic exchange with a main liquid phase in which macromolecules form a non-homogeneous solution.

Liquid-like phases have surface and bulk properties typical of liquids and acquire droplet morphology (29). Typical examples are the nucleolus, Cajal bodies, nuclear speckles, stress granules, P-bodies, and germ granules (30). Solid-like phases result from protein aggregation that causes microscopic solid deposition.

The morphology of solid deposits is exuberant. Crystal-like deposits are formed in endocrine and exocrine SGs (26, 28). Amyloids are ordered fibrillar protein deposits formed in neurodegenerative diseases, such as Alzheimer’s, Parkinson’s, Huntington’s, amyotrophic lateral sclerosis, Creutzfeldt–Jakob diseases, and type 2 diabetes (29–34) In the latter, islet amyloid polypeptide (IAPP) produced by pancreatic β-cells accumulates in the islets, leading to cell dysfunction, death, and insulin deficiency. On the other hand, typical amorphous aggregates in the eye lens cause cataracts, a widespread disease of aging.

Protein congregation involves heterotropic, non stoichiometric, protein–protein interactions, and it is characterized by its collective and cooperative nature. Also, it implies robust recognition mechanisms, acting simultaneously upon very different proteins. In this regard, it is revealing that proteins prone to congregation are frequently rich in intrinsically disordered regions (IDRs) and low complexity regions (LCRs), as if well-folded proteins were less susceptible to it. Partially unfolded protein conformations are ideal candidates for the promotion of collective congregation because they expose a higher proportion of active areas compared with native states, thus facilitating intermolecular over intramolecular interactions.

Given that cross-reactivity is an essential aspect of protein congregation, the identification and characterization of proteins that participate in congregates, and particularly of those that may display a congregating activity, promise to be of singular importance to boost our understanding of cellular processes, both in normal and pathological conditions.

In the present study we investigated in greater detail the biochemical and biophysical properties of ICA512 RESP18HD in relation to its congregating activity.

## Results

### ICA512 RESP18HD aggregates and binds Zn^2+^

The auto-aggregating activity of ICA512 RESP18HD, demonstrated by light scattering measurements (Fig. 2*A*), was negligible at pH 4.5, very slow at pH 6.8, and increased abruptly above pH 7.0.

**Fig. 2.**
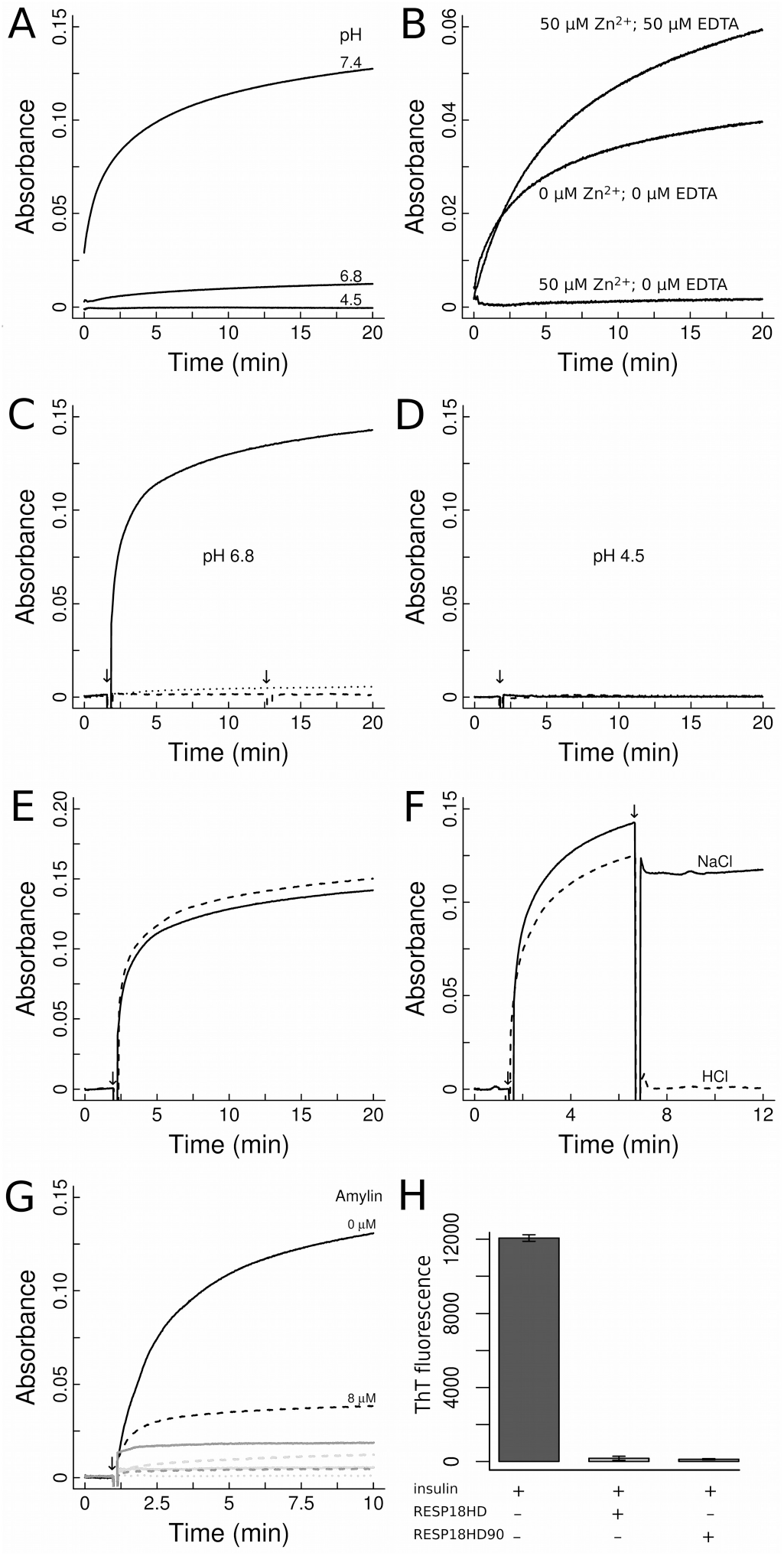
Time-course aggregation of ICA512 RESP18HD. The reaction was monitored by the increment of absorbance (ligth scattering) at 400 nm. In *A* and *B* the reaction was initiated adding ICA512 RESP18HD to the reaction buffer at 2 μM final concentration. In *C–G*, the reaction was initiated adding insulin 8 μM to 2 μM ICA512 RESP18HD solutions. Arrows mark the time of reagent addition. *A*: ICA512 RESP18HD aggregation is negligible at pH 4.5, barely detectable at pH 6.8, and very rapid at pH 7.4. *B*: Zn^2+^ inhibits aggregation of ICA512 RESP18HD at pH 7.4, an effect which is reversed by equimolar EDTA. *C*: incubation of ICA512 RESP18HD and insulin at pH 6.8 (solid line) results in aggregation. Insulin (dashed line) or ICA512 RESP18HD (dotted line) alone do not aggregate separately. The second arrow in the insulin trace marks the addition of 12.5 μM EDTA. *D*: neither insulin alone (dashes), ICA512 RESP18HD alone (dots), nor in combination (solid line) aggregate at pH 4.5. *E*: the aggregation reaction of insulin and ICA512 RESP18HD is similar in the presence or in the absence of 50 μM Zn^2+^ (dash and solid lines, respectively). *F*: aggregation at pH 6.8 is reversed by acidification to pH 4.0 with HCl (dash line; arrow at 400 s). Adding equivalent concentrations of NaCl instead of HCl stops further aggregation, but does not dissolve preexistent aggregates (solid line). *G*: amylin inhibits ICA512 RESP18HD–insulin aggregation. The reaction was performed in the absence of amylin (solid black line) and with 8 μM amylin (dashed black line). ICA512 RESP18HD is required for aggregation since neither 8 μM insulin + 8 μM amylin (solid gray line), 2 μM ICA512 RESP18HD + 8 μM amylin (dashed gray line), 8 μM amylin (solid light gray line), 2 μM ICA512 RESP18HD (dashed light gray line), nor 8 μM insulin (dotted light gray line) aggregate in the assay. *H*: ThT fluorescence assay for insulin fibrillation. Fibrillation was performed at 60 °C with stirring (see *Experimental Procedures*). The composition of the sample incubates is indicated at the bottom. Error bars indicate the standard deviation of three independent experiments.

This pH dependence suggested that ICA512 RESP18HD histidine residues (His 38, His 49 and/or His 98) might be involved in the aggregation. Therefore we tested possible histidine ligands. Micromolar concentrations of Zn^2+^ abolished aggregation, and the effect was reversible by equimolar EDTA (Fig. 2*B*). Neither Ca^2+^ nor Mg^2+^ acted like Zn^2+^, however Cu^2+^ mimicked Zn^2+^ but with lower efficiency (Fig. S1*A*).

Complete inhibition of ICA512 RESP18HD self-aggregation was observed at pH 7.4 and 15 μM Zn^2+^ (Fig. S1*B*). To roughly estimate ICA512 RESP18HD affinity for Zn^2+^, the aggregation kinetics was modeled as a first order reaction and an exponential decay was fit to the calculated rate constants (Fig. S1*C*). This simplified treatment allowed Zn^2+^ half-maximal inhibitory concentration to be estimated at 2 μM.

Direct equilibrium measurements of Zn^2+^ binding at pH 7.4 were complicated by aggregation. However, aggregation was minimal at pH 4.5 and microfiltration through 3 kDa membranes could be performed (not shown), and using the law of mass action a dissociation constant of 1 μM was calculated. This result suggests that ICA512 RESP18HD binds Zn^2+^ even in the acidic conditions of pH 5.5 present in the lumen of newly generated SGs (35), in which the concentration of Zn^2+^ is ~20 mM (36).

Far-UV CD spectra of ICA512 RESP18HD are shown in Fig. S1*D*. The spectrum of ICA512 RESP18HD at pH 4.5 corresponded to that of a largely unstructured peptide, and addition of 50 μM Zn^2+^ did not change significantly the spectrum. Both of these conditions inhibited the aggregation of ICA512 RESP18HD.

Instead, at pH 7.4 and in the presence of 50 μM Zn^2+^, a condition that also inhibits ICA512 RESP18HD aggregation, the CD spectrum evidenced the appearance of two negative bands at ~205 and ~218 nm, and one positive band at ~190 nm suggestive of α helix formation. At pH 7.4 and in the absence of 50 μM Zn^2+^ the spectrum taken at the onset of aggregation evidenced the same secondary structure signature. Based on these results, we hypothesize that fully unfolded ICA512 RESP18HD has little tendency to aggregate and that partial folding is required to achieve the aggregation-prone state.

To better delineate the ICA512 RESP18HD residues involved in the aggregation reaction, the variant 35–90 (RESP18HD90) was prepared. This variant contains the N-terminal Cys-rich motif (residues 35–61) but lacks the IDR encompassing residues 91–131 (see below). ICA512 RESP18HD90 did not aggregate under any of the conditions tested above (not shown).

### ICA512 RESP18HD–insulin aggregation

Unless otherwise indicated, the results reported in this section are from experiments conducted in incubation media containing 2 μM ICA512 RESP18HD and 8 μM insulin.

At pH 6.8, neither insulin nor ICA512 RESP18HD formed high-order aggregates in isolation. However, when incubated together, ICA512 RESP18HD and insulin formed aggregates, readily detectable by light scattering (Fig. 2*C*). ICA512 RESP18HD90 did not aggregate with insulin (not shown) and neither ICA512 RESP18HD nor insulin aggregated at pH 4.5 (Fig. 2*D*).

Zn^2+^ inhibited the self-aggregation of ICA512 RESP18HD at pH 7.4 (Fig. 2*B*), but it did not inhibit the coaggregation of insulin and ICA512 RESP18HD at pH 6.8 (Fig. 2*E*). Thus, the aggregation of ICA512 RESP18HD with insulin is not a consequence of previous aggregation of ICA512 RESP18HD.

To investigate if the observed aggregation of ICA512 RESP18HD with insulin at pH 6.8 could result from ICA512 RESP18HD removal of Zn^2+^ from insulin, it was verified that EDTA did not induce insulin aggregation (Fig. 2*C*).

Aggregates formed by coincubation of ICA512 RESP18HD and insulin at pH 6.8 dissolved at pH 4.0 (Fig. 2*F*). Moreover, aggregation was not influenced by reducing agents like DTT, phosphine or mercaptoethanol (not shown), suggesting that thiol–disulfide exchange reactions were not involved in the process.

ICA512 RESP18HD–insulin aggregation at pH 6.8 was completely blocked by preincubation with 100 mM l-arginine (not shown). l-arginine is an unspecific anti-aggregating agent employed in protein folding protocols (37). If l-arginine was added when aggregation was in progress, further aggregation was prevented but already formed aggregates did not dissolve.

ICA512 RESP18HD was also assayed for aggregation at pH 6.8 in the presence of proinsulin (6-His) and amylin. Proinsulin behaved similarly to insulin (not shown). Amylin was negative in the reaction with ICA512 RESP18HD but acted as a potent inhibitor in the aggregation of the pair ICA512 RESP18HD–insulin (Fig. 2*G*). This activity of amylin suggests that it interacts with either ICA512 RESP18HD, insulin, or with their complex blocking aggregation. Interestingly, an insulin anti-aggregating activity of amylin has been reported before (38).

In SGs, insulin is present mostly as a crystalline condensate and at concentrations as high as 30 mM (39). The concentration of soluble insulin potentially able to coaggregate with ICA512 RESP18HD is not known, but it could be higher than that assayed in the above experiments (8 μM). Therefore, the aggregation reaction was also tested at 2 μM ICA512 RESP18HD and 400 μM insulin. Aggregation was strongly inhibited under this condition (not shown).

The aggregate and soluble fractions from ICA512 RESP18HD–insulin incubates could be easily separated by centrifugation at 13,000 rpm. The recovery of significant amounts of both proteins in the pellet demonstrated that ICA512 RESP18HD and insulin coaggregate when incubated together at pH 6.8 (Fig. 3*A*). On the other hand, ICA512 RESP18HD90 did not coaggregate with insulin, as both proteins were not recovered together in the pellet (Fig. 3*A*).

**Fig. 3.**
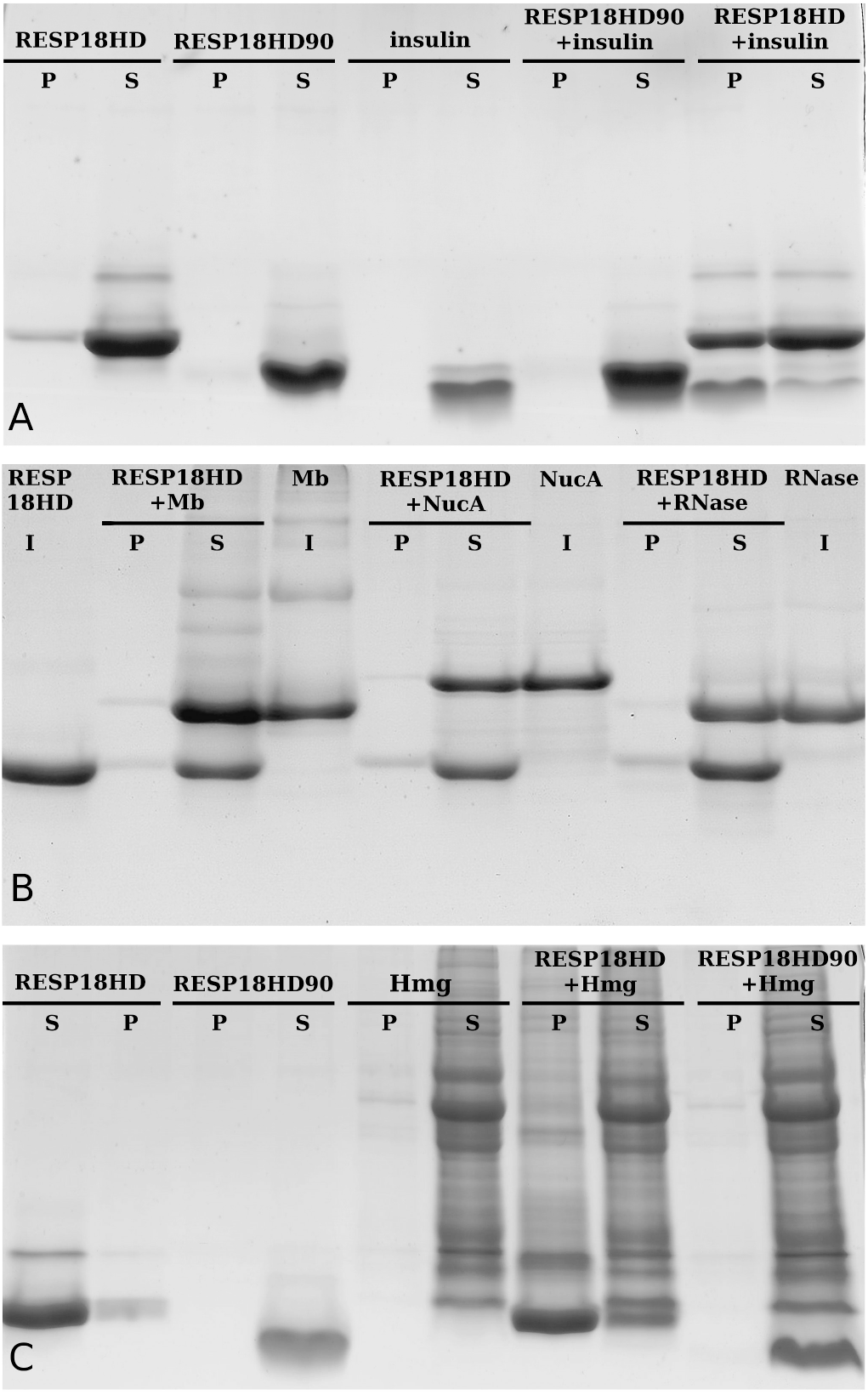
Isolation of ICA512 RESP18HD aggregates. The aggregation reaction was conducted as described in Fig. 2. Pellets (*P*) and supernatants (*S*) obtained by centrifugation from the sample incubates were loaded quantitatively in the SDS-PAGE gel lanes. *A*: co-incubation of insulin (8 μM) and ICA512 RESP18HD (2 μM) results in a pellet containing both proteins. ICA512 RESP18HD90 does not coaggregate with insulin and therefore both proteins are recovered only in the supernatant. Control lanes indicate that none of the proteins aggregate in isolation. *B*: ICA512 RESP18HD (2 μM) does not coaggregate with myoglobin, staphylococcal nuclease or RNase A (8 μM). These well-characterized, soluble proteins, along with ICA512 RESP18HD, are recovered in the soluble fractions (S). Centrifugation was omitted in control incubates (*I*). C: ICA512 RESP18HD (2 μM) exhibits a broad coaggregating activity upon a pancreas homogenate (Hmg). ICA512 RESP18HD90 lacks such activity.

Separation of pellets and supernatants followed by SDS-PAGE and band quantification established the proportion of each protein in the aggregate for different concentration ratios of ICA512 RESP18HD/insulin in the incubation media (Fig. *S2* and *S3*, and Methods *S1*). In excess of insulin, as is likely to occur in cells, the coincubation produced an aggregate with 2.8 moles of insulin per mole of ICA512 RESP18HD. In excess of ICA512 RESP18HD, the proportion was 0.4 mole of insulin per mole of ICA512 RESP18HD.

The variable proportion of each component suggests that the aggregate is formed by a solid-phase condensation mechanism, influenced by the relative concentrations of the interacting particles; which is different from specific mechanisms of aggregation involving constant stoichiometric relationships.

The effect of Zn^2+^ on the coaggregation reaction was further assessed by centrifugation and SDS-PAGE analysis (Fig. S4). This analysis indicated that at low concentrations Zn^2+^ does not inhibit the coaggregation, corroborating the above results obtained by light scattering measurements (Fig. 2*E*). However, 100 μM Zn^2+^ had a moderate inhibitory effect on the incorporation of ICA512 RESP18HD in the coaggregate.

### Temperature dependence of the aggregation

ICA512 RESP18HD self-aggregation and ICA512 RESP18HD–insulin coaggregation were tested at different temperatures in the 5–40 °C range. Aggregation velocities increased smoothly with temperature (not shown), as expected for typical collisional mechanisms. In a typical LLPT, for a given protein concentration, there is a critical temperature above which a single homogeneous phase exists (40). Such critical temperature could not be detected at the protein concentrations assayed in this work. The absence of a temperature-dependent cutoff in the aggregation reactions of ICA512 RESP18HD is compatible with solid-liquid phase separations, as those that result in microscopic deposits of amorphous aggregates or the formation of amyloid fibers.

Thioflavin T (ThT) is a highly specific probe for protein fibrillation because its fluorescence increases several orders of magnitude upon binding to the cross-β structure of amyloid fibrils. ICA512 RESP18HD self-aggregates and ICA512 RESP18HD–insulin coaggregates formed at 20 °C tested negative in a ThT fluorescence assay for cross-β fibrils (not shown), corroborating that the solid deposits were of amorphous nature.

### Inhibition of insulin fibrillation

Insulin submitted to high temperatures, stirring or agitation converts into amyloid cross–β fibrils (41). To test the possibility that ICA512 RESP18HD could affect the fibrillation of insulin, a fibrillation assay was set up in which insulin solutions were subjected to physical stress by incubation at 60 °C under stirring.

As expected, stressed insulin solutions yielded a strong ThT fluorescent signal. In the presence of either ICA512 RESP18HD or ICA512 RESP18HD90 ThT fluorescence was negligible, indicating that insulin fibrillation was abolished (Fig. 2*H*). Neither of the two ICA512 RESP18HD variants did fibrillate on their own account (not shown). These results imply that residues 35–90 of ICA512 RESP18HD bind to insulin and impede its fibrillation.

### ICA512 RESP18HD possesses broad protein condensing activity

To evaluate the specificity of the reaction, several pure, well-characterized and highly-soluble proteins were incubated with ICA512 RESP18HD under conditions similar to those used in the aggregation tests within insulin, and the pellets obtained by centrifugation were analyzed by SDS-PAGE. The results for myoglobin, staphylococcal nuclease, and RNase are shown in Fig. 3*B*. Similar results were obtained with lysozyme and human serum albumin (not shown). These results indicate that ICA512 RESP18HD has not condensing activity toward these isolated highly-soluble proteins.

Next, we asked if the condensing activity of ICA512 RESP18HD was preferentially directed to particular proteins in a complex mixture. To that end, the reaction was set up as above but replacing pure proteins by cell homogenates. The result with a rat pancreas homogenate is shown in Fig. 3*C*. In the absence of ICA512 RESP18HD, few proteins and in trace amounts were recovered in the pellet (Fig. 3*C*, *lane 5*). Adding ICA512 RESP18HD to the incubation mixture had a large effect in the reaction: nearly all proteins in the homogenate appeared with substantial yield in the pellet (compare lanes *7–8* with lanes 5–6 in Fig. 3*C*).

A density image analysis of homogenate coprecipitates illustrates the remarkable condensing activity of ICA512 RESP18HD (Fig. S5): in most cases at least half of each protein band was recovered in the pellet fraction, as well as nearly all ICA512 RESP18HD.

Interestingly, pure insulin added to the lysate was recovered in the pellet with low yield (Fig. S5). Thus, the affinity of ICA512 RESP18HD for insulin is counterbalanced by the competition for homogenate proteins that were in huge excess. Similar results were obtained with *S. cerevisiae* and liver homogenates (not shown).

Human EndoC-βH1 cells constitute an excellent experimental model to study insulin secretion (42). Therefore, the coaggregating activity of ICA512 RESP18HD was assayed with homogenates of these cells, and the results are shown in Fig. 4. ICA512 RESP18HD broadly coaggregated with EndoC-βH1 cell proteins (Fig. 4 lanes *3* and *4*), similarly to what was observed with other cell homogenates. Insulin *per se* did not coaggregate proteins in EndoC-βH1 cell homogenates (Fig. 4 lanes *11* and *12*). When in combination, ICA512 RESP18HD and insulin coaggregated along with the homogenate (Fig. 4 *lanes 5 and 10*). However, the yield of insulin in the coaggregate varied, with most insulin being recovered in the supernatant, as observed with rat pancreas homogenates (Fig. S5).

**Fig. 4.**
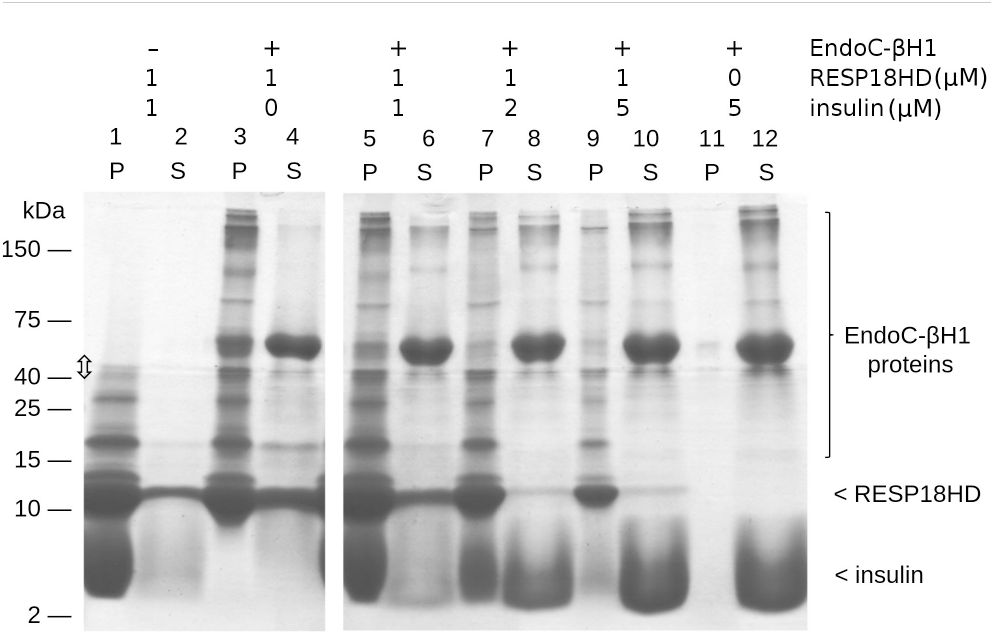
Coaggregation reaction of ICA512 RESP18HD and human EndoC-βH1 proteins. Two-layer SDS-PAGE separating gels were used in the analysis (10 and 17% acrylamide, top and bottom, respectively). The boundary between the layers is marked with a double-headed arrow. Fixed amounts of EndoC-βH1 homogenate were incubated with the indicated concentrations of ICA512 RESP18HD and insulin. The incubation mixture was processed as described in Fig. 3 to separate pellet (*P*) and soluble (*S*) fractions. *Lanes 1* and *2*: incubation of equimolar concentrations of insulin and ICA512 RESP18HD in the absence of EndoC-βH1 proteins results in coaggregation and recovery of most of both proteins in the pellet. Slow formation of covalently-linked multimers occurs spontaneously in ICA512 RESP18HD solutions (24). Here, however, this tendency was exacerbated by the coincubation with insulin. *Lanes 3* and *4*: ICA512 RESP18HD coaggregates with EndoC-βH1 proteins in the absence of added insulin. *Lanes 5–10: ICA512 RESP18HD* coaggregates with added insulin and EndoC-βH1 proteins. However, a five-fold excess of insulin over ICA512 RESP18HD decreases the yield of insulin in the coaggregate. *Lanes 11–12*: insulin *per se* has not aggregating activity on the protein homogenate.

### ICA512 RESP18HD–insulin condensates characterized by TEM

TEM analysis confirmed the amorphous nature of both ICA512 RESP18HD and ICA512 RESP18HD–insulin aggregates and the lack of fibers and regularly packed particles (Fig. 5). This result is coherent with the lack of fixed stoichiometry in the formation of the segregated solid phase described above. ICA512 RESP18HD–insulin preparations showed more dense aggregates than in the case of samples containing ICA512 RESP18HD alone.

**Fig. 5.**
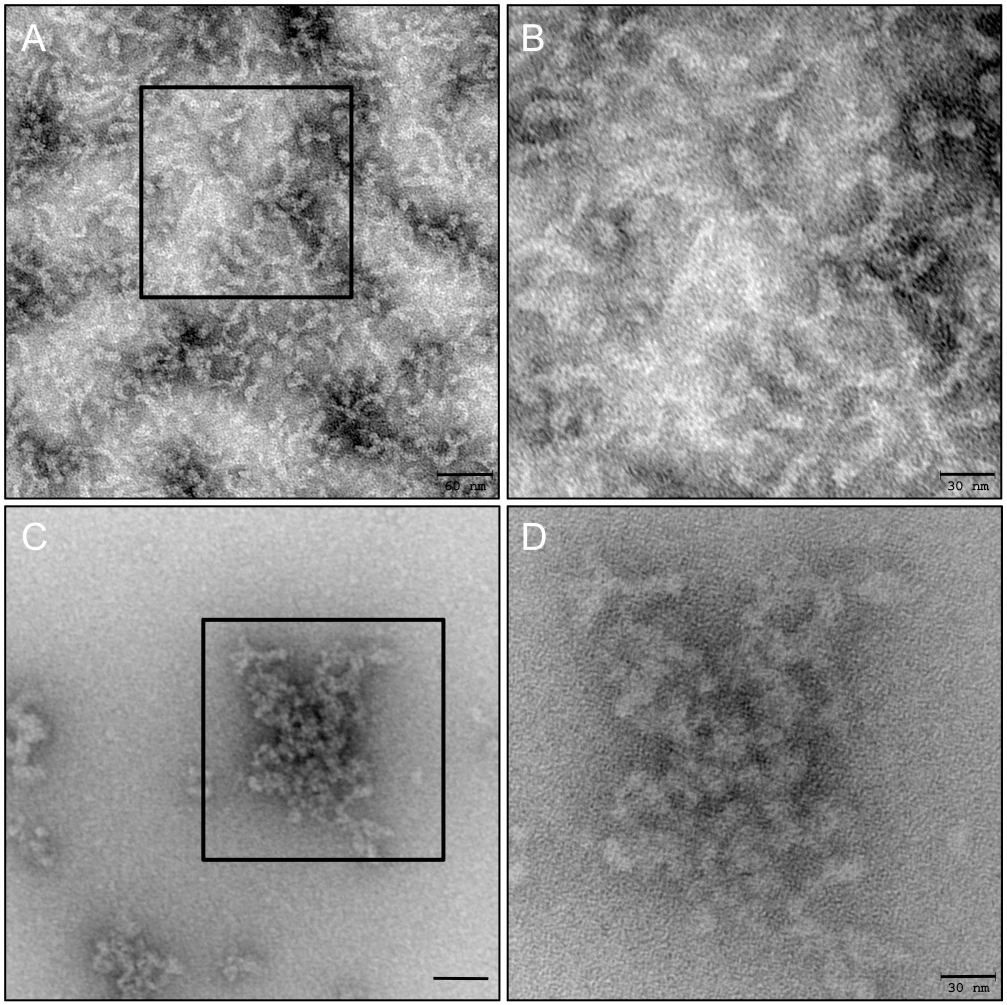
TEM of ICA512 RESP18HD and ICA512 RESP18HD–insulin aggregates. The aggregate of ICA512 RESP18HD and the coaggregate of ICA512 RESP18HD and insulin at pH 7.4 were prepared for negative staining. *A*: overview of ICA512 RESP18HD aggregates alone with higher magnification of the enclosed areas in *B. C* and *D*: low and high magnification images for ICA512 RESP18HD together with insulin. In both cases, the aggregate is of an amorphous nature and lacks periodic fibrils. The sponge-like matrix formed is thinner and more porous in the ICA512 RESP18HD alone aggregation. Scale bars: *A* and *C*: 60 nm, *B* and *D*: 30 nm.

### Residues 91–131 of ICA512 RESP18HD constitute an IDR

As shown above, ICA512 RESP18HD90 was unable to coaggregate with insulin. Therefore, it was inferred that the condensing activity of ICA512 RESP18HD requires residues 91–131. Using the sequence-based disorder prediction servers, PONDR VSL2 ((43); http://www.pondr.com/), IUPRED, and ANCHOR ((44); http://iupred2a.elte.hu/), residues 91–131 of ICA512 RESP18HD were identified as an IDR. This sequence region is enriched in arginine and proline residues (Fig. 1*B*), a typical signature of structural disorder.

### ICA512 RESP18HD antibody staining in cells and islets

Previously, we reported, that the C-terminally GFP-tagged ICA512 RESP18HD (residues 1–131) is targeted to insulin SGs, showing that ICA512 N-terminal domain contains sufficient information for SG localization in β-cell like INS-1 cells (20) r the current analyses, we transfected INS-1 cells with new constructs in which the GFP tag was replaced by its derivative Turquoise 2 (TQ2), TQ2 behaves essentially as GFP but is pH-insensitive and less prone to aggregation.

Also for these analyses, we generated a novel mouse monoclonal ICA512 RESP18HD antibody. Since ICA512 RESP18HD shares sequence homology with phogrin and Resp18, (Fig. 1*B*), we verified the specificity of this new antibody. To this end, INS-1 cells were transfected with human ICA512 RESP18HD–TQ2 (residues 1–131), Phogrin RESP18HD–TQ2 (residues 1–136) or Resp18–TQ2 (Resp18 residues 1–228) fusion proteins. INS-1 cells expressing the constructs were analyzed by immunoblotting, with either mouse anti-GFP (for the TQ2 tag) or the novel anti-ICA512 RESP18HD antibodies (Fig. 6*A*).

**Fig. 6.**
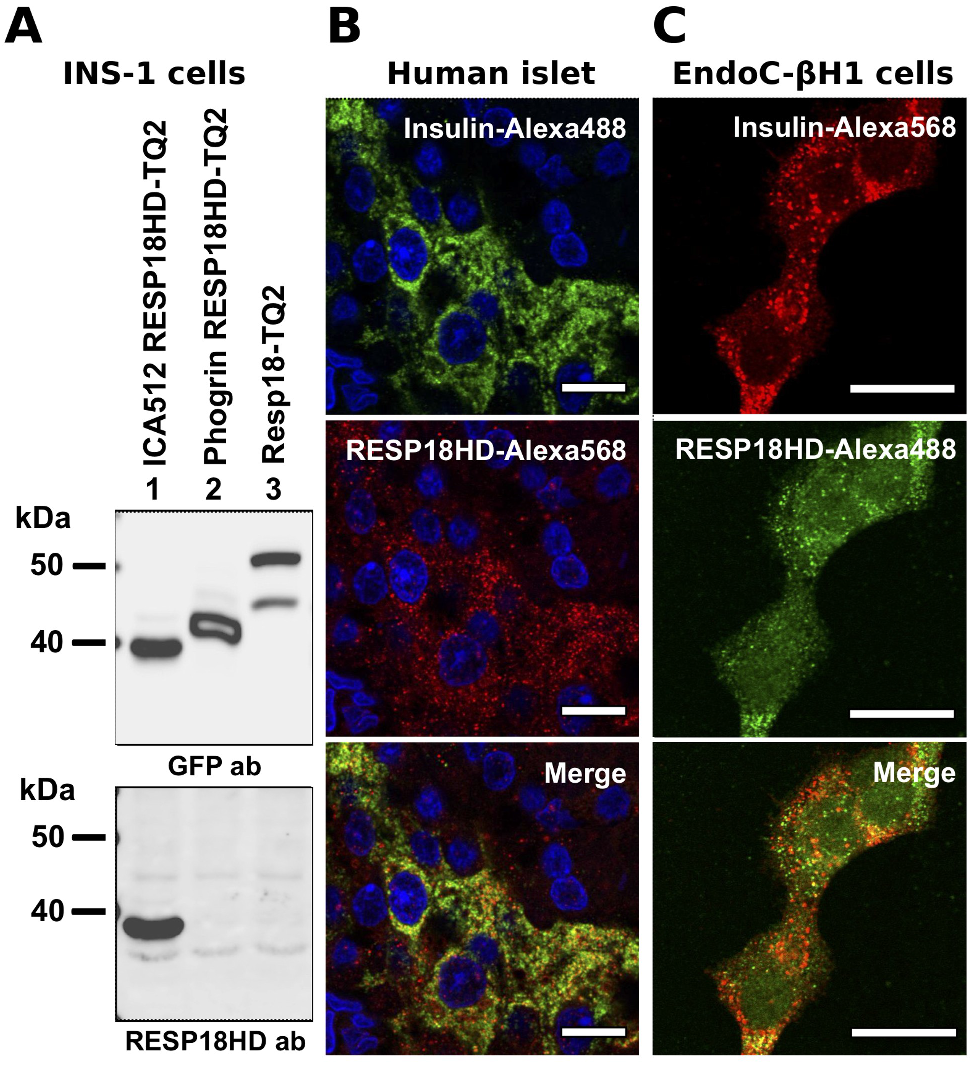
Immunostaining with the anti-ICA512 RESP18HD antibody. *A*: rat INS-1 cells transfected with constructs expressing C-terminal TQ2 fusion proteins of human ICA512 RESP18HD (residues 1–131), phogrin RESP18HD (residues 1–136) or Resp18 (residues 1–228) cells were lysed and analyzed by SDS-PAGE and immunoblotting, using for detection either mouse anti-GFP (that cross reacts with TQ2) or mouse anti-ICA512 RESP18HD antibodies. All intact TQ2 fusion proteins were detected by the anti-GFP antibody, whereas the anti-ICA512 RESP18HD antibody only recognized ICA512 RESP18HD–TQ2. *B*: immunostaining of human pancreatic tissue sections and *C*: human EndoC-βH1 cells with guinea pig anti-insulin and mouse anti-ICA512 RESP18HD antibodies. Insulin and ICA512 RESP18HD signals were found restricted to the islets with a punctuated signal consistent with SGs labeling (*B* and *C*). Scale bars indicate ~20 μm.

As expected, ICA512 RESP18HD–TQ2 was detected with both the anti-GFP and the anti-ICA512 RESP18HD antibodies, mainly as the intact fusion protein of ~40 kDa (Fig. 6*A*, lane *1*). Phogrin RESP18HD–TQ2 and Resp18–TQ2 were also recognized –mainly as intact fusions–by the anti-GFP antibody, but not by the anti-ICA512-RESP18HD antibody (Fig. 6*A*, *lanes* 2 and 3), demonstrating the specificity of the latter for ICA512. Further immunoblotting analysis using C-terminal deletion and sitespecific mutants established that the novel antibody recognizes a C-terminal epitope encompassing residue 125 of ICA512 RESP18HD (Fig. S6).

Next, the anti-ICA512 RESP18HD antibody was tested for detection of endogenous RESP18HD in human pancreatic tissue sections and human EndoC-βH1 cells by immuno fluorescence microscopy. The anti-ICA512 RESP18HD antibody signal was enriched in the endocrine cells and islets, where it overlapped with the signal for insulin (Fig. 6*B*). As expected, in EndoC-βH1 cells, the immunofluorescence signal for ICA512 RESP18HD colocalized with the punctuated insulin positive structures (Fig. 6*C*). Finally, the specificity of the anti ICA512 RESP18HD antibody was further corroborated by immunoblotting of human EndoC-βH1 cell lysates depleted of ICA512 by RNA interference (Fig. S7).

### ICA512 RESP18HD is processed upon regulated secretion

Next, we exploited the novel anti-ICA512 RESP18HD antibody to investigate the processing and putative secretion of ICA512 RESP18HD and variants thereof. To this aim, wild type and ICA512 RESP18HD–TQ2 transfected INS-1 cells were cultured in resting or glucose stimulated conditions, and the cell lysates and the immunoprecipitates from the corresponding culture media subjected to immunoblotting analysis (Fig. 7*A* and 7*B*, respectively).

**Fig. 7.**
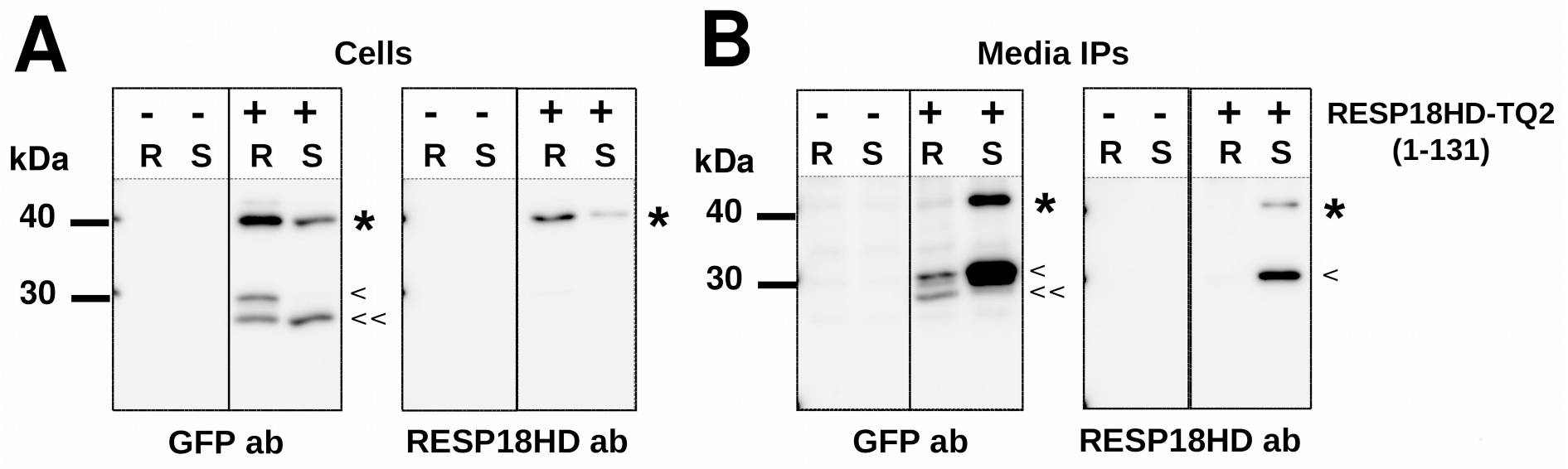
ICA512 RESP18HD–TQ2 processing in INS-1 cells. Resting (*R*) and glucose stimulated (*S*) conditions were assayed. Immunoblotting was performed using mouse anti-GFP or anti-ICA512 RESP18HD antibodies. *A*: cell lysates. *B*: culture media proteins immunoprecipitated with a goat anti-GFP antibody. The amount of ICA512 RESP18HD–TQ2 (~40 kDa; *asterisks*) was reduced in stimulated compared to resting cells. The opposite applied to media immunoprecipitates. This behavior is consistent with regulated secretion of ICA512 RESP18HD–TQ2. Remarkably, a single fragment of ~30 kDa (*arrowhead*) comprising a short C-terminal sequence of ICA512 RESP18HD preceding TQ2 was highly enriched in the culture media from stimulated cells, implying that the proteolysis of the intact 40 kDa fragment is coupled to its secretion. Other proteolytic fragments of ~27 kDa (*double arrowheads*) did not react with the anti-ICA512 RESP18HD antibody.

In resting and stimulated INS-1 cells, full-length ‘mature’ ICA512 RESP18HD–TQ2 was detected by anti-GFP and anti-ICA512 RESP18HD antibodies as a band of ~40 kDa (Fig. 7*A* *asterisks*). In resting cells the anti-GFP antibody detected also a faint band >40 kDa, indicating the presence of proICA512 RESP18HD–TQ2 species still containing the signal peptide.

Mature ICA512 RESP18HD–TQ2 detected with both antibodies was more abundant in resting than in stimulated cells, which is compatible with its secretion upon glucose stimulation.

In resting, but not in stimulated cells, the anti-GFP antibody recognized an ICA512 RESP18HD–TQ2 proteolytic fragment of ~30 kDa (Fig. 7*A* *arrowhead*). This fragment corresponds to TQ2 preceded by a short sequence from the C-terminus of ICA512–RESP18HD. Its absence in stimulated cells suggests that it may be exported along with the mature form upon stimulation.

Another proteolytic fragment of ~27 kDa compatible with the bare size of TQ2 was detected in resting and stimulated cells (Fig. 7*A* *double arrowheads*). The fact that this fragment is not depleted in stimulated cells is an indication that it may be not secreted, as often observed for cytosolic GFP/TQ2 remaining after incomplete proteasomal degradation of fusion proteins ((45) and references therein).

Remarkably, in the culture media, the 40 kDa, mature ICA512 RESP18HD–TQ2 was present in significant amounts only with stimulated cells (Fig. 7*B* *asterisks*), which were also enriched for the presence of a single ~30 kDa fragment (Fig. 7*B* *arrowhead*). The abundant recovery of both ICA512 RESP18HD–TQ2 species in the culture media of glucose stimulated cells suggests that the 40 kDa fragment is secreted in a regulated fashion and converted to the 30 kDa species by proteolysis.

### ICA512 RESP18HD processing site

Immunoprecipitates from culture media of glucose stimulated INS-1 cells expressing ICA512 RESP18HD–TQ2 were run on SDS-PAGE, and the isolated ~30 kDa band subjected to ‘in gel’ digestion with trypsin and mass spectroscopy analysis.

Observed peptide masses and fragmentation patterns (not shown) identified (a) several peptides corresponding to the inner sequence of TQ2 and (b) two N-terminally extended peptides with the sequence ‘drsglapgpvatmvsk’ and ‘sglapgpvatmvsk’, respectively. In these two peptides, ‘drsglap’ and ‘sglap’ map to the C-terminus of ICA512 RESP18HD (see also Fig. 1*B*; residues 125–131, and 127–131, respectively), ‘gpvat’ corresponds to a linker sequence between ICA512 RESP18HD and TQ2, and ‘mvsk’ is from the N-terminus of TQ2. The shorter peptide is likely result of the trypsin digestion applied. Since the single ~30 kDa fragment reacted with the anti-ICA512 RESP18HD antibody (Fig. 7*A* *arrowhead*), that detects an epitope including the residue 125, (see also Fig. S6) it is concluded that the processing site locates close to residue 125 (see Fig. 1*B*) and within the IDR of ICA512 RESP18HD. Due to limitations of the ICA512 RESP18HD antibody epitope recognition and also other potential peptide recognition by the mass spectroscopy analysis, we cannot completely exclude the presence of other proteolytic sites beyond D125.

## Discussion

Our finding that ICA512 RESP18HD possesses *in vitro* protein condensing activity turns it into a potentially important factor for SG biogenesis and regulated secretion. In this regard, ICA512 RESP18HD resembles other lumenal domains of SG membrane proteins, including dopamine β-hydroxylase, peptidyl glycine α-amidating enzyme, carboxypeptidase E, and granin, which have been reported to coaggregate with SG cargoes proteins (Ref. (46) and references therein). Interestingly, these condensing factors aggregate at the acidic pH of mature SGs whereas ICA512 RESP18HD displays condensing activity at pH close to neutral, with acidic pH inhibiting its self-aggregation or coaggregation with insulin. Hence ICA512 RESP18HD likely exerts its condensing activity at early stages of granulogenesis along the secretory pathway, *i.e*. prior to that of other granule cargoes.

The coaggregation of ICA512 RESP18HD and insulin takes place at very low protein concentrations, under pH and temperature conditions close to physiological, and with no external physical stress (shearing, stirring, or heating). Under these conditions, isolated ICA512 RESP18HD and insulin *per se* do not aggregate appreciably and exhibit a moderate and low degree of oligomerization, respectively (24, 47). Thus, proper conditions for coaggregation of insulin and ICA512 RESP18HD are likely to evolve *in vivo*.

Another important finding is that Zn^2+^ does not inhibit ICA512 RESP18HD–insulin coaggregation (Fig. 2*E*). Since Zn^2+^ inhibits the homophylic aggregation of ICA512 RESP18HD at neutral pH (Fig. 2*B*), the lack of effect on the coaggregation with insulin suggests that the reaction is not triggered by previously formed aggregates of ICA512 RESP18HD and, on the contrary, is caused by the interaction of soluble ICA512 RESP18HD and insulin. This behavior also suggests that coaggregation would not be impeded by the high concentrations of Zn^2+^ present in the SG of β–cells. On the other hand, the coaggregation of ICA512 RESP18HD and insulin may be resolved upon their exposure to the very acid pH of mature SGs (Fig. 2*D* and 2*F*). Moreover, the observation that amylin inhibits the coaggregation of ICA512 RESP18HD and insulin (Fig. 2*G*), reinforces the idea of an interaction between two soluble partners that results in mutual insolubility, for amylin is known to interact directly with soluble insulin and impair its aggregation (38).

Several stable, soluble, and well-characterized proteins were shown to coaggregate if incubated at their melting temperature. This indicates that partially folded states are involved in coaggregation (48). ICA512 RESP18HD is known to be partially unfolded under our experimental conditions. Insulin, on the other hand, is monomeric and folded. However, monomeric insulin is less stable than hexameric insulin and has a greater tendency to populate partially folded states (49). Thus, if coaggregation of ICA512 RESP18HD and insulin requires two partially unfolded partners, ICA512 RESP18HD might act shifting the equilibrium of monomeric insulin toward a partially folded state.

Our findings buttresses the idea that one of the main mechanisms involved in targeting and regulated secretion is the congregation of mutually interacting proteins in aggregated phases (26, 28). However, due to the complexity of the machinery for granule biogenesis, the multiplicity of players, and their redundancy, an essential role of ICA512 RESP18HD is unlikely. Proof of which is that in mice with genetic deletion of ICA512 and phogrin, two proteins with a RESP18 domain, the size of SG stores are reduced but not abolished (6).

Coaggregation of ICA512 RESP18HD–insulin *in vitro* occurs within a few minutes and under very mild conditions, and therefore is different from typical insulin fibrillation or amyloidogenic reactions, which take days and require physical stress. However, partially folded states and aggregates can evolve towards highly stable cross-β fibrils, and this process is at the core several misfolding diseases and attracts enormous interest from a practical and theoretical point of view (50).

For its availability in pure form and large quantities, and its tendency to form cross-β structure *in vitro*, insulin is a popular experimental model for the study of fibrillogenesis. Even though insulin fibrils have not been directly linked to any misfolding disease, they have been associated with problems in the manufacturing, storing, and administration of insulin (50, 51). In addition, cytotoxic effects of insulin fibrils have been reported in cell cultures (52), opening the possibility that yet to be discovered misfolding disorders directly due to insulin fibrillation might exist.

Predictably, biophysical studies found that insulin fibrillogenesis has much in common with the amyloidogenesis of several proteins associated with misfolding pathologies. The coincidences arise because fibrillation and amyloidogenesis are manifestations of protein folding, a fundamental process in biology. For fibrillation to occur, conformational changes must take place, such us the backbone adoption of β–structure and its association into regular β–sheets perpendicular to the fibril long axis. All proteins, to a greater or lesser degree, can form fibrillar structures, because for this folding process to occur, the sequence details are much less important than for proper folding into the complex and elaborated native state. Indeed, the regularity of the peptide bond determines the regularity of the fibers, as opposite to the native state, for which specific side chain contacts are crucial determinants for the overall fold of the backbone.

The very general nature of the fibrillation process allows many instances for intervention, and thus, fibrillation can be inhibited in multiple different ways. For instance, chemical and physical agents that stabilize the native state inhibit fibrillation. Natural and synthetic ligands act in the same direction by displacing the equilibrium toward the native state.

Ligands interacting with the fibrillation nucleus or with protofibrils may block the formation of cross-β structures. Driven by the need of therapeutic agents for amyloidosis, several chemical compounds, peptides, and proteins have been discovered with the potential to stop the progression or even revert fibrillation processes (53).

The discovery in this work that the interaction between ICA512 RESP18HD and insulin results in amorphous coaggregation prompted us to test if ICA512 RESP18HD itself would fibrillate and induce the fibrillation of insulin.

Unexpectedly, not only ICA512 RESP18HD was unable to fibrillate itself but was also a strong inhibitor of the fibrillation of insulin. These effects were recapitulated with ICA512 RESP18HD90, suggesting that the N-terminal segment of ICA512 RESP18HD is responsible for this inhibition. Since ICA512 RESP18HD90 lacks the C-terminal IDR and does not aggregate *per se*, its previous aggregation is dispensable for inhibition of insulin fibrillation.

Two previously proposed mechanisms for the inhibition of insulin fibrillation are (a) binding of the inhibitor to a fibrillation competent partially folded insulin monomer, and (b) direct binding of the inhibitor to the growing fibrils (49). Further studies are needed to define which of these alternatives apply to ICA512 RESP18HD inhibition.

Insulin fibrils, or precursors thereof, have been proposed to affect β-cells and insulin secretion and circulation. Thus presumably an efficient control mechanism keeps in check insulin fibrillation. This presumption is further sustained by the fact that, despite the high concentrations of insulin in β-cells and the easiness of insulin fibrillation *in vitro*, no pathologies directly associated to insulin fibrillation have ever been reported.

A very strong inhibitor of insulin fibrillation could be the very high intragranular concentration of insulin; which, along with the high concentration of Zn^2+^, stabilizes the hexameric insulin–Zn complex and reduces the concentration of aggregation-prone monomeric insulin.

Amylin, which is present at high concentrations in the SG, may also be a natural inhibitor of insulin aggregation and fibrillogenesis (54, 55). Also an ubiquitous Ca^2+^-binding protein, nucleobindin–1, similar to calreticulin, was found to inhibit insulin fibrillation, attenuate fibril-induced cell toxicity, and proposed to act as an insulin chaperone in the control of amyloidogenesis in type 2 diabetes (56).

Moreover, we found that exocytosis of ICA512 RESP18HD is coupled with the proteolysis of its C-terminal IDR. Since this is the region responsible for the condensing activity of ICA512 RESP18HD at pH >7.0, it is tempting to propose that such cleavage is required for preventing detrimental consequences resulting from the aggregation of the protein upon its secretion in the extracellular compartment. The identification of the protease responsible for ICA512 RESP18 HD IDR cleavage and the regulation of this process shall be the topic of future studies.

In conclusion, given the multiplicity of proteins and peptides that interact in the SG, it can be hypothesized that the control mechanism to prevent amyloidogenesis is based in a web of interactions between granular cargo and constitutive proteins. In the context of this complex anti amyloidogenesis system, ICA512 RESP18HD may play an important role due to its preferential interaction with insulin. However, new *in vivo* experimental approaches need to be devised to confirm this possibility.

## Experimental procedures

### Antibodies, chemicals and general protocols

Chemicals were of the purest analytical grade available. Human insulin was a gift from Dr. Nestor Annibali (Denver Farma, Buenos Aires, Argentina) and also purchased from Sigma (91077C). Recombinant human proinsulin extended with N-terminal Met, 6-His tag and Lys was from R&D Systems (1336-PN). Thioflavin T was from Sigma (T3516).

The novel mouse monoclonal antibody against human ICA512 RESP18HD was raised with the MPI–CBG antibody facility (Dresden, Germany). Mouse and goat anti-GFP, as well as guinea pig anti-insulin antibodies were described previously (20).

In some experiments, to achieve good resolution in a wide range of molecular weights, SDS-PAGE gels prepared with two consecutive separating layers of 10% and 17% acrylamide, respectively.

ICA512 RESP18HD concentration was measured using an absorption coefficient at 280 nm of 7,450 M^−1^ cm^−1^ (24). Binding of Zn^2+^ to ICA512 RESP18HD was assessed by filtration through microfilters of 3 kDa cutoff. Zn^2+^ concentration was determined colorimetrically, following the formation of a complex with Zincon (SC-25839 Santa Cruz Biotechnology).

CD spectra were collected at 20 °C on a Jasco 810 spectropolarimeter (Jasco Corporation, Tokyo, Japan) as described (57). Steady-state fluorescence measurements were performed on an ISS K2 multifrequency phase fluorometer (ISS, Champaign, Illinois, USA) equipped with a cell holder connected to a circulating water bath at 20 °C and with 1.0-cm cells.

In-gel trypsin digestion of SDS-PAGE excised bands was performed with an In-Gel Tryptic Digestion Kit (Thermo Scientific). Recovered peptides were subjected to LC-MS/MS with electrospray ionization in a LTQ Orbitrap XL ETD instrument (Thermo Scientific).

Microscopy images of human EndoC-βH1 cells and pancreatic islet sections were acquired with an Olympus FluoView-1000 laser scanning confocal microscope equipped with a 60 PlanoApo OLSM lens (numerical aperture 1.10) (MPI-CBG, Dresden, Germany). Image analysis was carried out with ImageJ (https://imagej.nih.gov).

### DNA constructs and protein expression

The pET-9 plasmid for expression of human ICA512 RESP18HD in *E. coli* was previously described (24). A truncated variant ICA512 RESP18HD90 (residues 35–90) was generated by site-directed mutagenesis (QuikChange, Stratagene) by replacing the codon for residue 91 with a stop codon. Purification of ICA512 RESP18HD was previously described (24). A similar protocol was used to purify ICA512 RESP18HD90.

For expression in INS-1 cells, cDNAs encoding ICA512 N-terminal signal peptide sequence and either full (residues 1–131) or a truncated variant of ICA512 RESP18HD (residues 1–124) was generated by PCR and fused in frame to a sequence encoding Turquoise2 (TQ2) at their C-termini. TQ2 is a derivative of GFP, and ICA512 RESP18HD–GFP was assessed in our previous work (20). Similarly the cDNAs encoding for both human phogrin ICA512 RESP18HD (residues 1–136) and human Resp18 (residue 1–228) were generated by PCR and fused at their C-termini to TQ2. All constructs were verified by DNA sequencing.

### Cell culture and immunocytochemistry

Culture of INS-1 insulinoma cells, their exposure to resting (R) and high glucose stimulation (S) of the cells, cell lysate preparation, and culture media immunoprecipitation (IP) were performed as described (20). To harvest R and S culture media, these were at first pipetted from the cells into 15 ml Falcon tubes, inclusive protease inhibitors (1:100; Sigma), centrifuged briefly to clear the media fractions from any residual cells, and prepared for immunoprecipitation (IP) using goat anti-GFP antibody essentially as described in (20). Human EndoC-βH1 cells were cultured in DMEM low glucose (1 g/l), 2% BSA Fraction V, 50 μM 2-mercaptoethanol, 10 mM nicotinamide, 5.5 μg/ml transferrin, 6.7 ng/ml selenite, 100 U/ml penicillin and 100 μg/ml streptomycin. EndoC-βH1 cell lysates were prepared in 50 mM HEPES, pH 7.0. Paraffin sections of human pancreatic islets were prepared as described (58). EndoC-βH1 cells and the pancreatic islet sections were immunostained with guinea-pig anti-insulin and the novel mouse anti-ICA512 RESP18HD antibody and the respective Alexa-conjugated secondary antibodies, as described previously (20, 58).

### Aggregation reactions

Time-course of ICA512 RESP18HD aggregation at 20 °C was followed measuring scattered light as UV absorption at 400 nm. The stock solution of ICA512 RESP18HD was prepared in 25 mM sodium acetate, pH 4.5. The reaction was started adding ICA512 RESP18HD to a final concentration of 2 μM to 25 mM HEPES adjusted to the indicated pH in the 4.5–7.4 range by adding acetic acid or sodium hydroxide. To test the effect of different metals and EDTA, these were added to the reaction buffer and the pH readjusted to the desired value before starting the measurements.

Coaggregation of ICA512 RESP18HD with insulin was assayed by light scattering as indicated above. The reaction was started adding insulin to ICA512 RESP18HD solutions, resulting in 8 and 2 μM final concentration, respectively. Different additives to the basic reaction were as described in *Results*.

### ThT fluorescence assay

To induce the formation of insulin amyloid cross-β fibrils, 20 μM insulin in 100 mM HEPES, pH 6.8 was incubated for 1 h at 60 °C with a stirring bar rotating at 150 rpm. At time zero and at the end of the incubation, samples were withdraw, diluted to 2 μM, and added to 2 μM ThT in the same buffer for immediate fluorescence measurement. Excitation and emission wavelength were 440 and 480 nm, respectively. ICA512 RESP18HD and ICA512 RESP18HD–insulin aggregates formed by incubation at 20 °C were directly assayed for ThT fluorescence without heating and stirring treatment.

### Separation of aggregate proteins by centrifugation and SDS-PAGE analysis

The formation of high-order protein aggregates was analyzed by centrifugation and SDS-PAGE. Typically, 10 μM ICA512 RESP18HD with or without other proteins at similar concentration was incubated for 30 min in 100 μL of 100 mM HEPES, pH 6.8. The samples were then spun for 5 min at 13,000 rpm in a microcentrifuge. The supernatant and pellet fractions were TCA-precipitated and pellets dissolved in SDS-PAGE sample buffer. Samples were heated for 5 min at 60 °C in the presence of 100 mM DTT before loading into the SDS-PAGE gel.

### Transmission electron microscopy (TEM)

ICA512 RESP18HD aggregates, prepared as described in the previous section after incubation alone or together with insulin, were pipetted onto mesh grids equipped with a carbon film and incubated for 4 min. Afterwards excess fluid was removed and the grids were incubated with 1% uranlylformate solution for 30 sec. After removal of most of the staining solution the grids were air-dryed. TEM images were acquired with a Tecnai 12 Biotwin transmission electron microscope (FEI Company) operated at 120 kV equipped with a bottom-mount 2×2K F214 CCD camera (TVIPS).

## Acknowledgments

We thank Daniela Richter for the inmmunostaining of human pancreatic tissue sections; Jürgen Weitz, Marius Distler, Nicole Radisch for the provision of surgical pancreatic specimens; Patrick Keller for antibody production at MPI-CBG; and Marc Gentzel (Biotec, Dresden) for mass spectroscopy analysis, and Katja Pfriem for outstanding administrative assistance.

## Author contributions

MS and MRE conceived the study, designed the experiments and supervised the project; PLT expressed and purified recombinant proteins, performed biophysical studies; JMT designed constructs, performed molecular studies in cultured cells; AM carried out the EM studies; AS was responsible for cell culture and transfection; CW was responsible for molecular cloning; MS and MRE interpreted the data and wrote the paper with contributions from PLT, JMT and AM.

## The abbreviations used are

RPTP: receptor-type protein tyrosine phosphatase
ICA512: islet cell antigen 512
SG: secretory granule
PTP: protein tyrosine phosphatase
NTF: N-terminal fragment
TMF: transmembrane fragment
MPE: membrane proximal ectodomain
Resp18: regulated endocrine-specific protein 18
RESP18HD: RESP18 homology domain
LLPT: liquid-liquid phase transition
IDR: intrinsically disordered region
IAPP: islet amyloid polypeptide
ThT: Thioflavin T
TQ2: Turquoise 2
CD: circular dichroism

