## Supplemental Figs S1-S7 and Methods S1 for "ICA512 RESP18 homology domain is a protein condensing factor and insulin fibrillation inhibitor"

### List of the supplementary data

**Methods S1.** Stoichiometry of ICA512 RESP18HD–insulin aggregates
**Fig. S1.** Aggregation of ICA512 RESP18HD and metal binding
**Fig. S2.** Concentration dependence of the formation of high-order ICA512 RESP18HD–insulin aggregates analyzed by centrifugation and SDS–PAGE
**Fig. S3.** Insulin and ICA512 RESP18HD composition in the aggregate
**Fig. S4.** $Zn^{2+}$ at high concentration diminishes ICA512 RESP18HD–insulin coaggregation
**Fig. S5.** Coaggregation of ICA512 RESP18HD and pancreas homogenate proteins
**Fig. S6.** ICA512 RESP18HD antibody recognizes an epitope encompassing residue D125
**Fig. S7.** Detection of ICA512/IA-2/PTPRN using the novel anti-ICA512 RESP18HD antibody and RNAi mediated knockdown in human EndoC–βH1 cells

### Methods S1. Stoichiometry of RESP18HD–insulin aggregates

To investigate the proportion of of each protein in the aggregate, 1—20 µM ICA512 RESP18HD was incubated with 10 µM insulin for 30 min in 100 µL of 100 mM HEPES, pH 6.8. Then the samples were centrifuged 5 min at 13000 rpm using a microcentrifuge. The supernatant (*S*) and pellet (*P*) fractions were TCA-precipitated and the TCA pellets redissolved in SDS–PAGE sample buffer. Samples were incubated 5 min at 60 °C in the presence of 100 mM DTT and analyzed by SDS-PAGE gel with Coomasie blue staining (Fig. *S2*).

Bands in the gels shown in Fig. *S2* were quantified using *ImageJ (https://imagej.nih.gov)*. The integrated intensity value for each band was used to calculate the fraction of each protein that is incorporated into the pellet and the molar ratio of both components in the pellet.

A numeric example will clarify the procedure for the calculations. Take the lanes labeled 7.5:10 in Fig. S2*A*, which corresponds to 7.5 µM of ICA512 RESP18HD and 10 µM insulin in the incubation media. For *P* and *S*, the integrated intensity of the RESP18HD band is $i_P$ and $i_S$, respectively. Thus, the total intensity ($i_T$) is $i_P + i_S$. Because RESP18HD is either in *P* or in *S*, $i_T$ represents the total amount of ICA512 RESP18HD in the incubation. The fraction of ICA512 RESP18HD in *P* and *S* is calculated as

$$f_P = i_P / i_T, \text{ and } f_S = i_S / i_T.$$

The fraction of insulin in *P* and *S* is calculated in the same way.

Fractions calculations are always performed with intensities from the same protein and therefore there is no need to consider differences in the intrinsic staining capacity. In the example, $f_P$=0.73 and the moles of RESP18HD in the pellet will be $n_{RESP18HD}=0.73 \times 7.5 \times 10^{-6}$ M $\times v$, where $v$ is the incubation volume. Moles of insulin in the pellet can be calculated in the same way:

$$f_P = 0.77 \text{ and } n_{insulin} \times 7.5 \times 10^{-6} \text{ M} \times v$$

Finally, the ratio ICA512 RESP18HD/insulin in the aggregate can be calculated as

$$r = n_{RESP18HD}/n_{insulin}$$

The results of the calculations are shown in Fig. *S3* below.

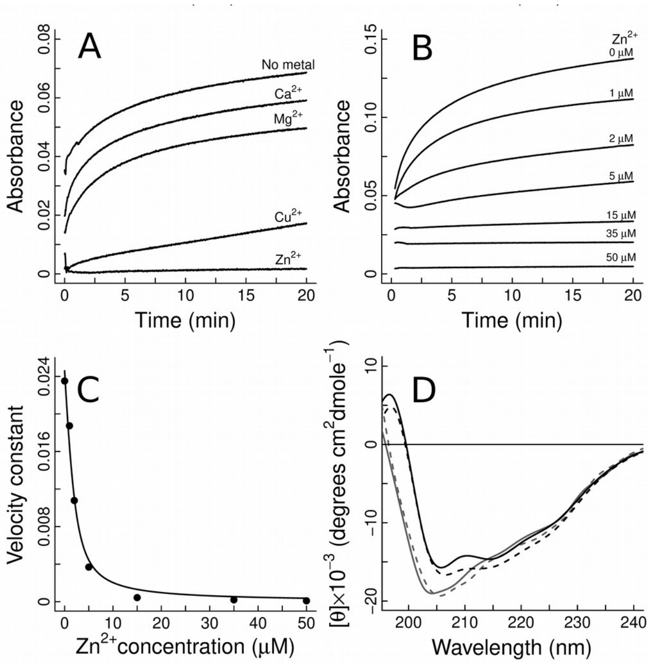


**Fig. S1.** Aggregation of ICA512 RESP18HD and metal binding. The time-course of ICA512 RESP18HD aggregation was followed measuring scattered light as UV absorption at 400 nm. The reaction buffer was HEPES 25 mM, pH 7.4, containing no added cations, $Zn^{2+}$, $Ca^{2+}$, $Mg^{2+}$, or $Cu^{2+}$ at the indicated concentrations. The reaction was started adding ICA512 RESP18HD to a final concentration of 2 µM. *A*: at 50 µM concentration, neither $Ca^{2+}$ nor $Mg^{2+}$ inhibit ICA512 RESP18HD aggregation; however $Cu^{2+}$ does it, albeit with lower efficiency than $Zn^{2+}$. *B*: $Zn^{2+}$ inhibits aggregation in a concentration-dependent manner. Onset and nearly complete of inhibition occur at sub micromolar and 15 µM $Zn^{2+}$ concentration, respectively. *C*: The aggregation kinetics was modeled as a first-order reaction, and the calculated rate constants were represented as a function of $Zn^{2+}$ concentration. The half-maximal inhibitory concentration estimated from the fit curve was 2 µM. *D*: Far UV-CD spectra show that ICA512 RESP18HD is largely unstructured at pH 4.5, whether in the presence of 50 µM $Zn^{2+}$ or not (gray dashed and full lines, respectively). At pH 7.4 and without added $Zn^{2+}$, a condition that promotes ICA512 RESP18HD aggregation, the appearance of two negative bands at ∼205 and ∼218 nm and one positive band at ∼190 nm denotes an increase in α helix content (black line). At pH 7.4 and with 50 µM $Zn^{2+}$, a condition that inhibits aggregation, structuring is somewhat accentuated (black dashed line).

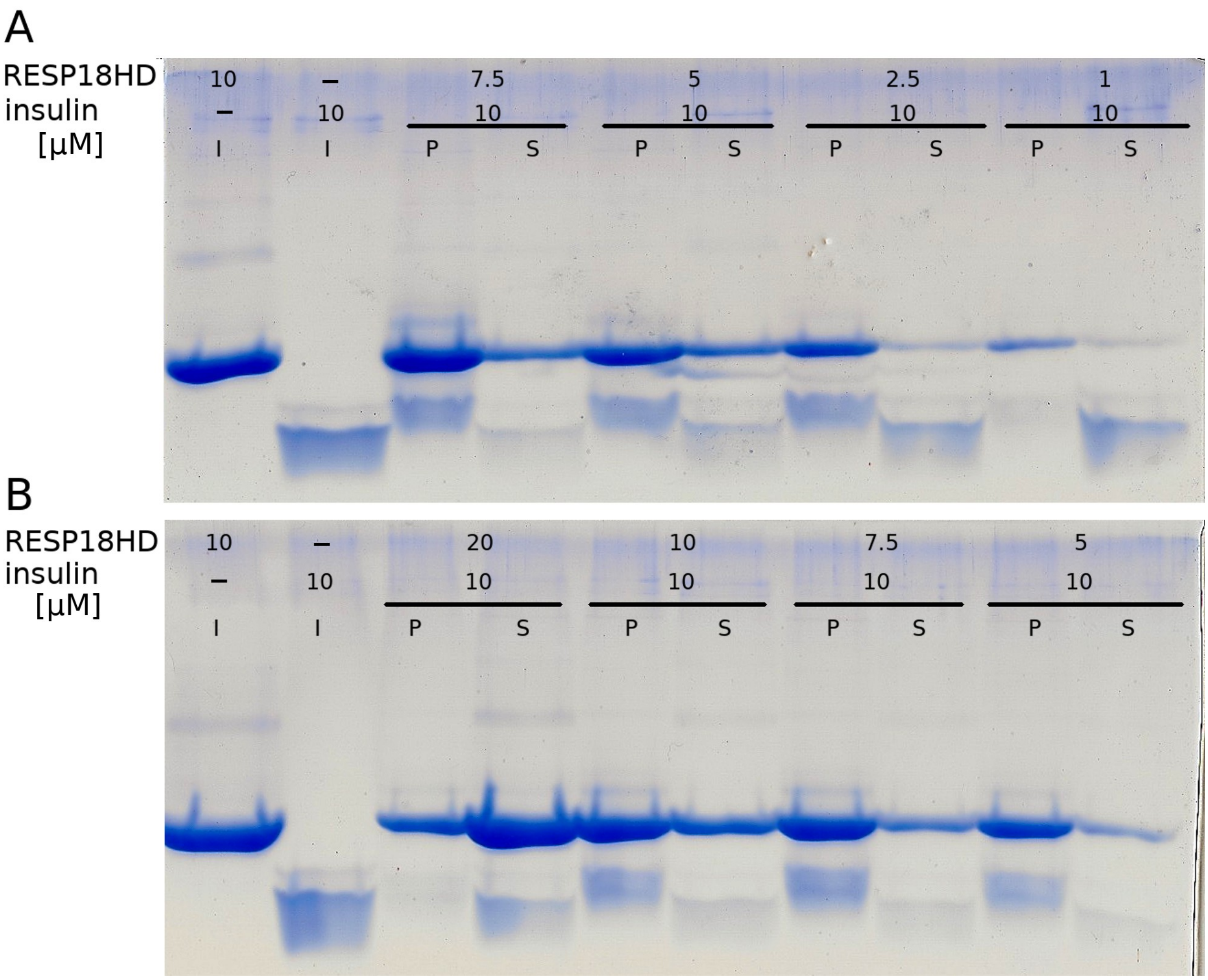


**Fig. S2.** Concentration dependence of the formation of high-order ICA512 RESP18HD–insulin aggregates analyzed by centrifugation and SDS–PAGE. *A* and *B*: Numbers indicate the concentration (µM) of each protein in the incubation media. Pellet, supernatant, and incubate are labeled *P*, *S*, and *I*, respectively.

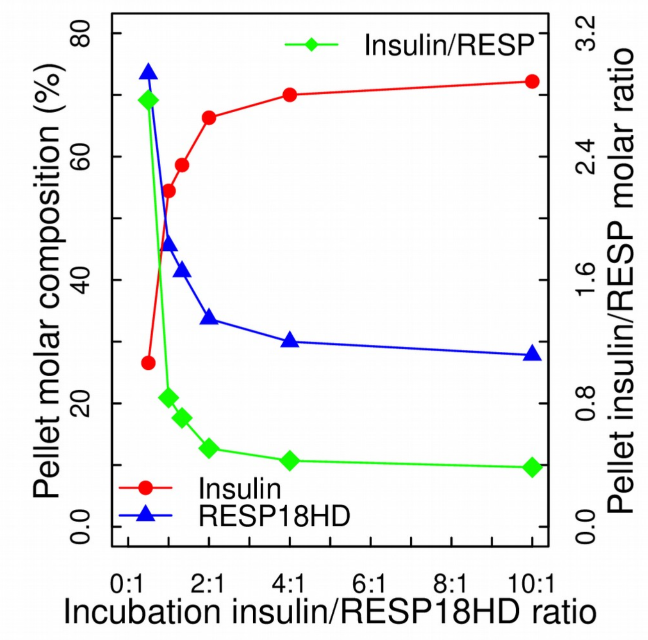


**Fig. S3.** Insulin and ICA512 RESP18HD composition in the aggregate. The fraction of each incubated protein recovered in the pellet for the experiment in Fig. S2 is represented as a function of insulin/ICA512 RESP18HD molar ratio in the incubation media (insulin, red; ICA512 RESP18HD, blue; left axis). The insulin/ICA512 RESP18HD ratio in the aggregate is also shown (green, right axis). The amount of each protein in pellets and supernatants was determined by image analysis of SDS–PAGE of gels in Fig. S2. The concentration of insulin in the sample incubates was 10 μM; the concentration of ICA512 RESP18HD varied between 1 and 20 μM for each experimental point. Using the formulae described in Methods S1, it can be calculated that at low concentrations, ICA512 RESP18HD condensates very efficiently insulin. In these conditions, the aggregates contain on average about 2.8 molecules of insulin per molecule of ICA512 RESP18HD. At high concentrations, ICA512 RESP18HD is less efficient: most insulin and ICA512 RESP18HD remain in solution and the aggregate contains about 0.4 molecules of insulin per molecule of ICA512 RESP18HD.

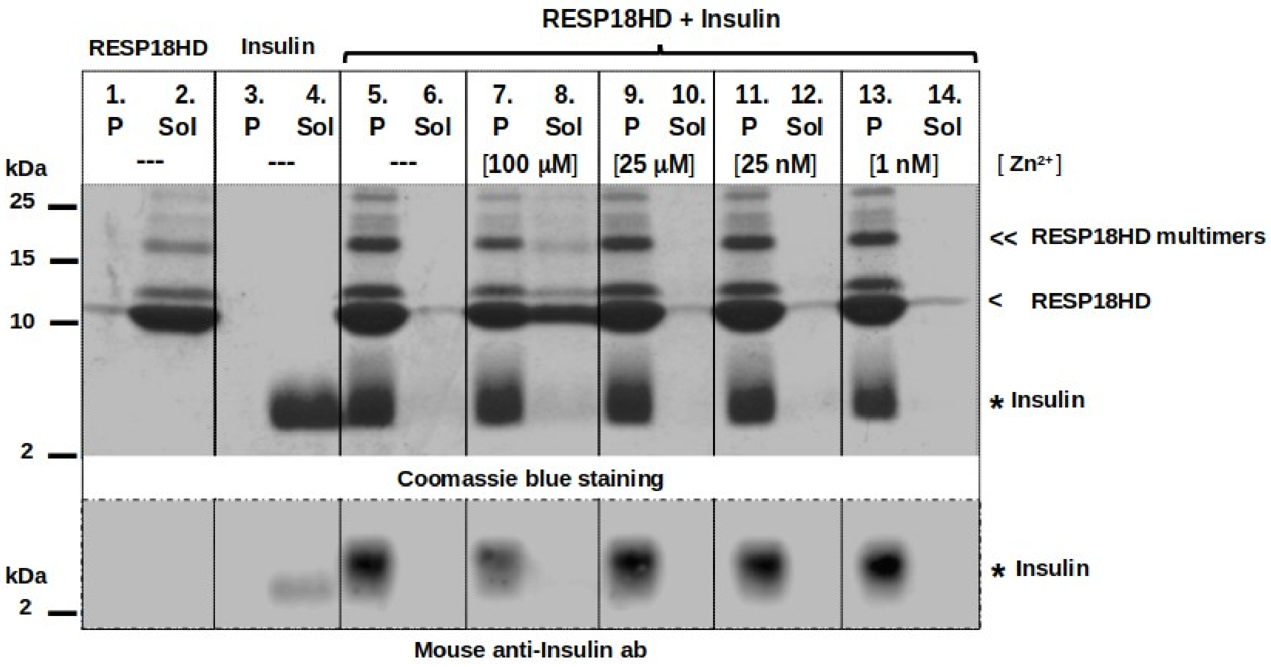


**Fig. S4.** $Zn^{2+}$ at high concentration diminishes ICA512 RESP18HD–insulin coaggregation. The coaggregation reaction and separation of aggregate (*P*) and soluble fractions (*S*) were performed as described in the main text of the article. 10 µM ICA512 RESP18HD, 10 µM insulin, and $Zn^{2+}$ were included in the reaction as indicated. 100 µM $Zn^{2+}$ exerts a moderate inhibitory effect on the reaction. This inhibition affects ICA512 RESP18HD incorporation in the pellet without affecting insulin aggregation. The formation of soluble multimers of ICA512 RESP18HD during storage and incubation has been described before (Sosa et al. 2016, Biochim. Biophys. Acta, 1864 511–522). Here, it is shown that the formation of ICA512 RESP18HD multimers is exacerbated by the incubation with insulin.

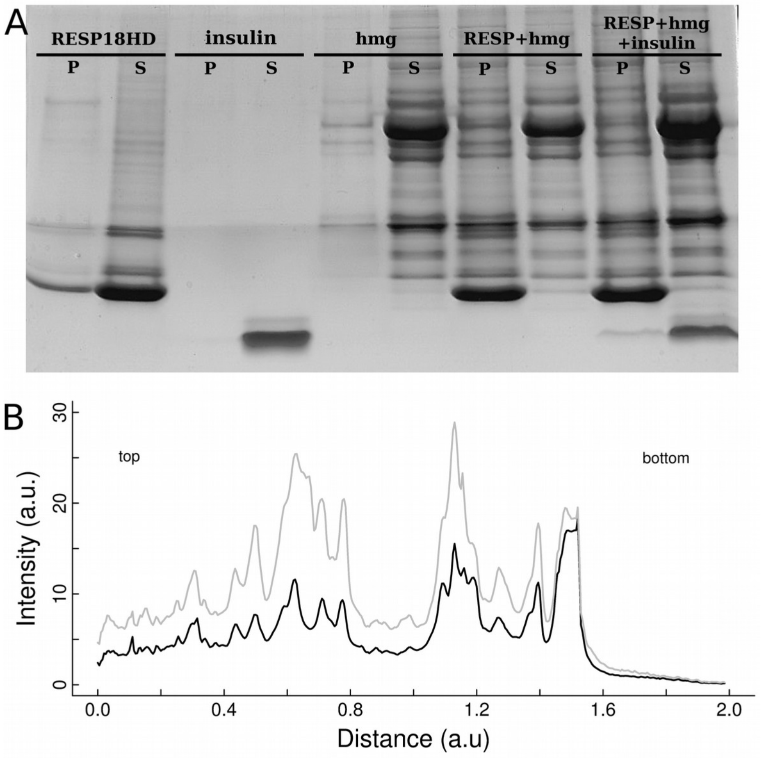


**Fig. S5.** Coaggregation of ICA512 RESP18HD and pancreas homogenate proteins. The coaggregate isolated by centrifugation was analyzed by SDS–PAGE, and Coomassie blue stained bands were quantified with *ImageJ* (see the Experimental Procedures in the main text for the details). *A*: aggregation of a rat pancreas homogenate (*hmg*). Comparison of *7th* and *8th* lanes shows that most of the homogenate proteins incubated with ICA512 RESP18HD are recovered with good yield in the pellet (*P*) separated by centrifugation from the soluble protein (*S*). *B*: *7th* and *8th* lanes in *A*, were analyzed to calculate the intensity of the stained bands along the SDS–PAGE gel. The gray line represents the sum of the intensities of the two *lanes* (*P* + *S*). The black line represents the intensity of *7th lane* (*P*).

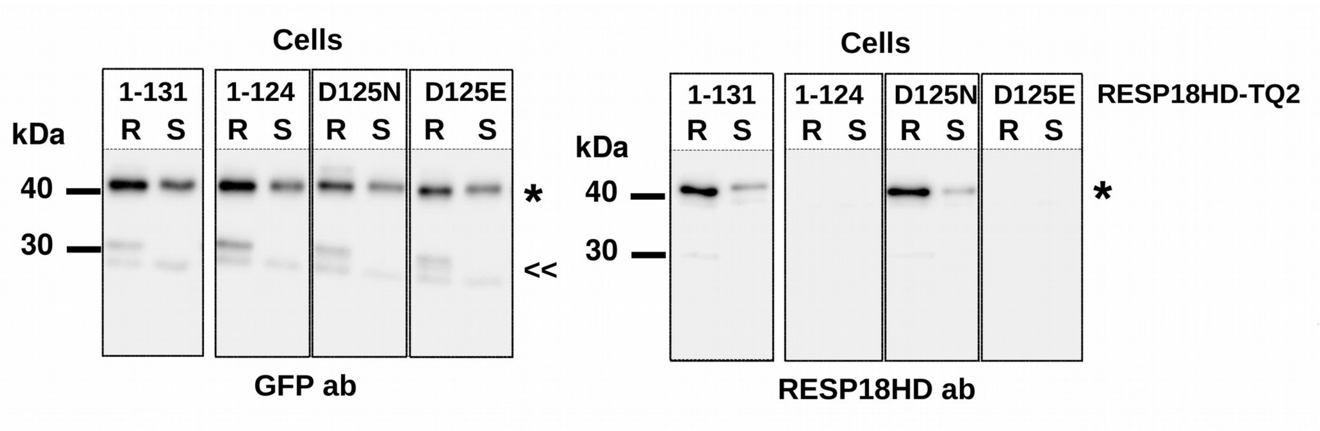


**Fig. S6.** ICA512 RESP18HD antibody recognizes an epitope encompassing residue D125. Lysates from resting (*R*) and glucose stimulated (*S*) INS-1 cells expressing full length ICA512 RESP18HD–TQ2 (residues 1–131), a truncated mutant (residues 1–124) and point mutants D125N and D125E of the full length variant (residues 1–131) were analyzed by immunoblotting using mouse anti-ICA512 RESP18HD and mouse anti-GFP antibodies. Mature ICA512 RESP18HD–TQ2 (*asterisk*) and proteolytical fragments thereof (arrowheads) were recognized by the anti-GFP antibody. Notably, anti-ICA512 RESP18HD antibody recognized the mature form of 1-131 and its D125N mutant, but did not recognize the truncated form of 1–124 or the D125E mutant. Thus, anti-ICA512 RESP18HD antibody recognizes an epitope including D125.

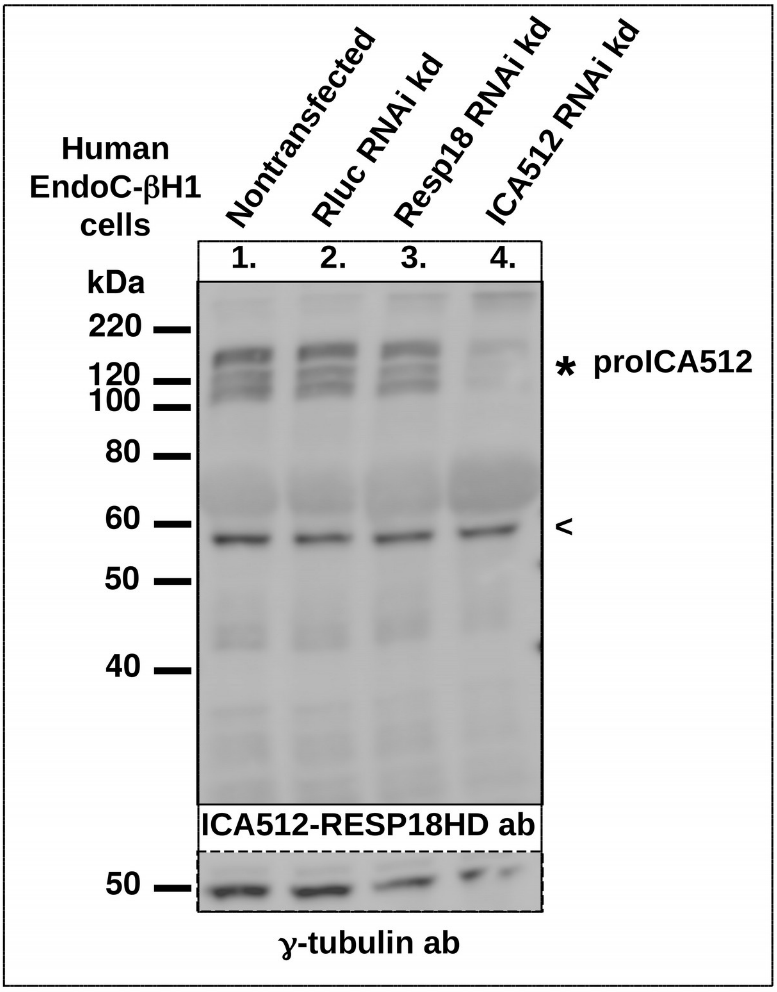


**Fig. S7.** Detection of ICA512 using the novel anti-ICA512 RESP18HD antibody and RNAi mediated knockdown in human EndoC–βH1 cells. Renilla lucilla (Rluc), human Resp18, or human ICA512 specific 'esiRNA' (Eupheria Biotech, Dresden) RNAi was used for transfection of human EndoC–βH1 cells. Cells were lysed three days post transfection and analyzed for the influence of the respective RNAi's by immunoblotting and using the novel antibody raised against human ICA512 RESP18HD (MPI-CBG and PLID, Dresden, Germany). As expected, the antibody detected multiple fragments ≥ 100 kDa (*asterisk*), corresponding to non- and to variedly glycosylated species of proICA512. The arrowhead indicates a nonspecific protein, conceivably heavy chain IgG from the cell culture medium, recognized by the secondary anti-IgG antibody. Pronounced depletion of the detected proICA512 species was apparent upon treatment of the cells with ICA512 specific RNAi (lane *4*), whereas no apparent effect on ICA512 expression was observed upon Rluc or Resp18 RNAi treatments (lanes *2* and *3*).
